# Interpretable and Accurate Prediction Models for Metagenomics Data

**DOI:** 10.1101/409144

**Authors:** Edi Prifti, Yann Chevaleyre, Blaise Hanczar, Eugeni Belda, Antoine Danchin, Karine Clément, Jean-Daniel Zucker

## Abstract

Biomarker discovery using metagenomic data is becoming more prevalent for patient diagnosis, prognosis and risk evaluation. Selected groups of microbial features provide signatures that characterize host disease states such as cancer or cardio-metabolic diseases. Yet, the current predictive models stemming from machine learning still behave as black boxes. Moreover, they seldom generalize well when learned on small datasets. Here, we introduce an original approach that focuses on three models inspired by microbial ecosystem interactions: the addition, subtraction, and ratio of microbial taxon abundances. While being extremely simple, their performance is surprisingly good and compares to or is better than Random Forest, SVM or Elastic Net. Such models besides being interpretable, allow distilling biological information of the predictive core-variables. Collectively, this approach builds up both reliable and trustworthy diagnostic decisions while agreeing with societal and legal pressure that require explainable AI models in the medical domain.

## INTRODUCTION

An increasing wealth of data from high-throughput molecular and imaging technologies is connecting medical sciences and machine learning (ML). The latter is impacting numerous areas of medicine, including disease diagnosis and prognosis ^1-3^. It is now argued that deep learning, a field of ML, will become the most beneficial technology to hit radiology since digital imaging and that ML will dramatically improve prognosis within the coming years ^4^.

Simultaneously, progress made in high throughput technologies has contributed to developing new fields such as metagenomics, which allows qualifying and quantifying microbial ecosystem composition and functionality with unprecedented resolution. The association of the gut microbiota with human health and disease has been widely discussed ^5^ and links with numerous diseases such as obesity^6^, liver cirrhosis ^7^, type I ^8^ and type 2 diabetes ^9^, inflammatory bowel disease ^10^, and colorectal cancer ^11^ have been described. Although these associations are proposed as predictive, many of these findings are only correlative and require controlling for confounding factors. This task remains a challenging objective ^12^.

Ecological relationships among bacterial species such as mutualism, parasitism, and competition ^13^ may change with a shift in microbial equilibrium. Although causality is challenging to establish, identifying easily interpretable markers of microbial shifts can allow predicting disease states and/or progression. Some authors accurately predicted low microbial richness individuals ^14^ and we confirmed these predictors in independent populations ^15^. Others discriminated liver cirrhosis patients from controls using metagenomes ^7^. Such metagenomics predictors were also proposed in other conditions such as obesity, type-2 diabetes, IBD, and colorectal cancer ^9,11,16,17^.

Despite these findings, metagenomics data must be analysed carefully because they are often performed in a small number of samples (*N*) compared to a very large number of variables (*p*). Current microbial catalogues, which are composed of millions of genes ^18^ and thousands of bacterial species and functional profiles ^19^, allow characterizing and comparing sampled ecosystems. As a consequence, most models tend to overfit the training data and result in predictions arising from random sampling fluctuations. To reduce overfitting, some authors use learning algorithms that include a dimension reduction or regularization methods, e.g. Elastic Net ^11^ or SVM-RFE ^12^. While these algorithms are more straightforward than others, they generate complex models that are difficult to interpret. ML research has focused on building accurate models for large data collections, often at the expense of interpretability.

Providing an *explanation* of the prediction process is increasingly requested in precision medicine, especially before validating and deploying the model in patient care ^20^. In Europe, the new GDPR legislation defines that explanation of prediction models is a necessity ^21^. The *comprehensibility* — the extent to which a human can make sense of a model — is not necessarily sufficient to ensure that the model is validated. Ideally, experts need *justifiability*, defined as being in line with existing domain knowledge. Interpretable models have two desirable properties: conciseness and readability by nonexperts. They should contain simple operations (e.g. addition using integers) and be limited in size ^22^. Some authors consider the sparse linear models produced by the Lasso algorithm as interpretable ^23^. For others, the models should be presented as a decision tree or a list of rules. Tibshirani introduced the “sparsity bet” claiming that if the “true model” was complex, then we would need much more data than what are available to learn it accurately. As a consequence, learning a sparse approximation (i.e. small number of features) is the *best* one can do ^24^.

Collectively, these aspects of a model defined as interpretability, are at the core of the present work. Causality, as the holy grail of modern biology, is out of the scope of the interpretability property of a predictive model. Here, we hypothesized that models inspired by ecosystem relationships and sparse microbial signatures can be accurate and more interpretable than state-of-the-art (i.e. SOTA) models, including logistic regression with elastic-net regularization (ENET) and support vector machines (SVM).

## RESULTS

### A new family of models for metagenomics data

We propose a new family of models, named BTR for Binary/Ternary/Ratio, which are an oversimplification of linear models. For each ecosystem *y*_1_… *y*_n_, the abundance or presence of either genes, taxonomy levels, functions, or other microbial qualities is represented by *X*_1_… *X*_p_ predictor variables.A patient is predicted in a disease state with a probability of 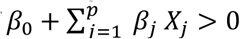where *β*_0_…*β*_p_ are the real coefficients of a linear model. The biological assumptions are that the contribution of each bacterial species to the prediction is proportional to its abundance and that only a limited number of species is sufficient to support the prediction. BTR models are much simpler and improve interpretability without worsening accuracy. Our models are inspired by three hypotheses emphasizing relationships between species and associated ecosystem outcomes (**Figure 1**).

**Figure 1:**
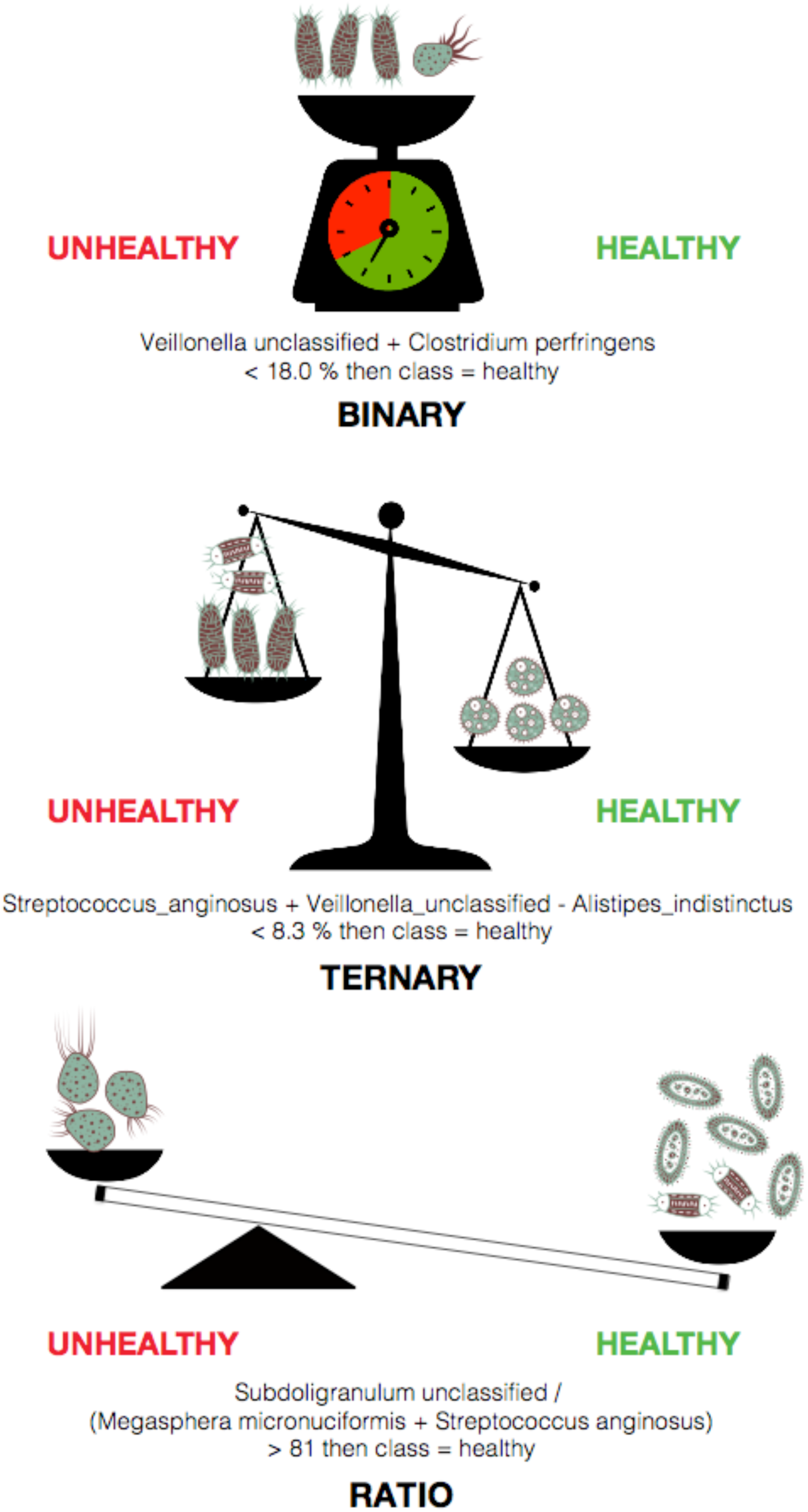
The three balance concepts depicting the BTR models. *Top*: The Binary model tests whether the cumulated abundance of a set of species is below or above a certain threshold. *Middle*: The Ternary model tests whether the cumulated abundance of a first set of species is below or above the cumulated abundance of a second set of species plus a certain threshold. *Bottom*: The Ratio model tests whether the cumulated abundance of a first set of species over the cumulated abundance of a second set of species is above a given threshold.

Hypothesis 1: The unweighted cumulative abundance of a group of species can predict disease state. We define the *binary models* (*i.e.* Bin) as linear models with the additional constraint that each coefficient *β*_1_ *β*_*p*_ (omitting the intercept *β*_0._) must be binary — 0 or 1 (**Figure 3A**; see online methods; *(1)*). Biologically, these species may not interact directly with each other (*e.g*. non-overlapping resources or are not co-located) or be associated together (*e.g*. cooperation, or similar ecological niche ^25,26^). Hypothesis 2: The difference of unweighted cumulative abundance of two groups of species can predict disease state. This assumption is implemented by *ternary models* (*i.e.* Ter). These are also linear models with the constraint that each coefficient *β*_*1*_… *β*_*p*_ (omitting the intercept *β*_0._) is limited to the value −1, 0 or 1 (**Figure 3B**; see online methods; *(2)*). Hypothesis 3: The ratio of unweighted cumulative abundance of two groups of species can predict disease state. This assumption is implemented by *ratio models* (*i.e.* Ratio), which are also linear models with an additional constraint: each coefficient *β*_*1*_ *β*_*p*_ is limited to a value of *-θ*, 0 or 1, where *θ* is a positive real number, and the intercept *β*_*0*_ is set to zero (**Figure 3C**; see online methods; *(3))*. Biologically, both Ter and Ratio models can correspond to different types of species interactions including simultaneous cooperation and competition between species.

**Figure 3:**
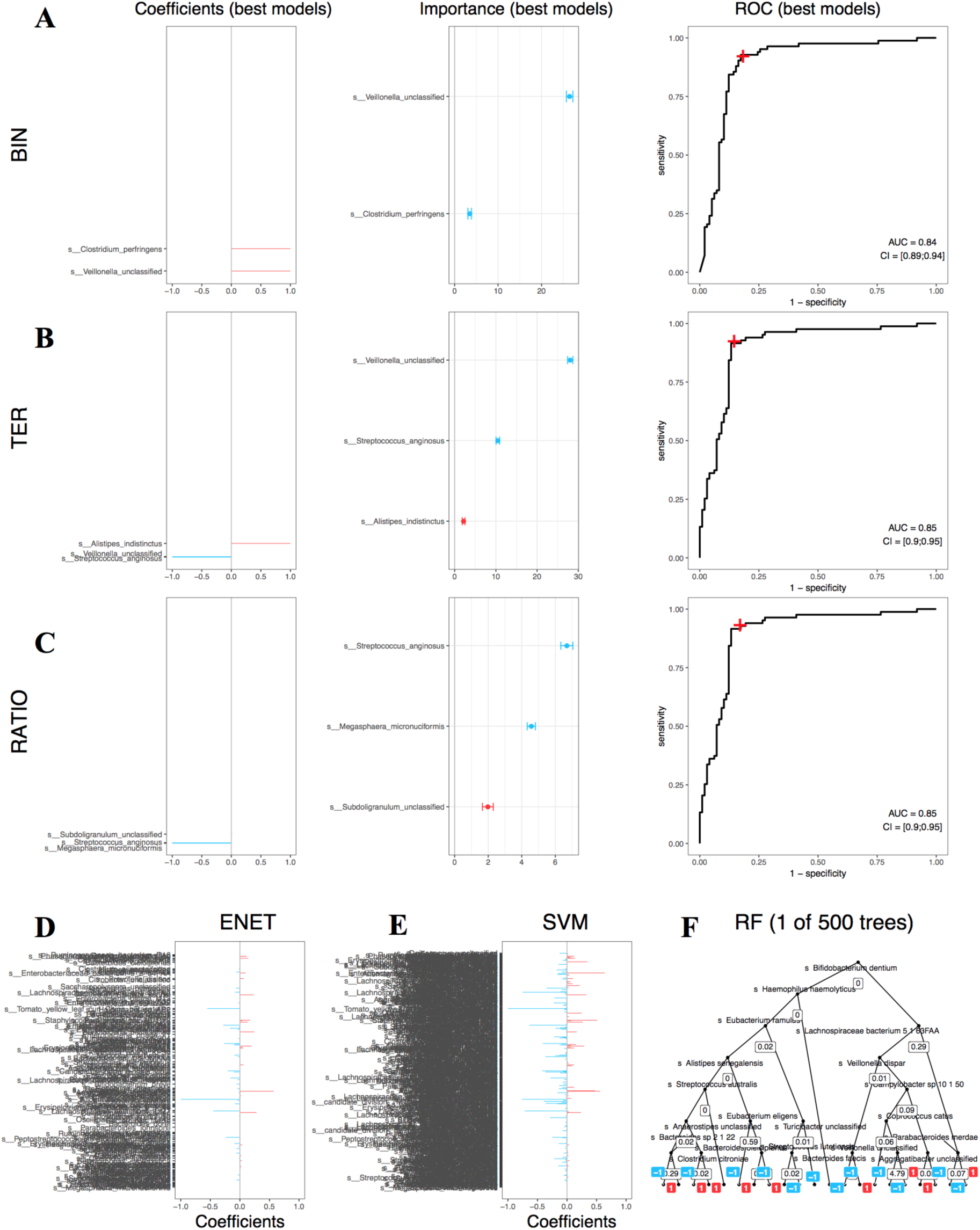
BTR models are interpretable compared to state-of-the-art. ***A-C*** *left*: Barcode graphical representations indicating the coefficients of the BTR model features sorted by decreased correlation strength with the class to predict. **A-C** *middle*: Mean decrease accuracy (MDA) plots indicating feature importance computed during the cross-validation process. Blue and red colours indicate enrichment in patients and controls respectively. **A-C** *right*: Receiver operator characteristic (ROC) plots for the same BTR models. The red cross indicates the specificity and sensitivity of the model. **D-F**: A visualisation attempt of the SOTA models with barcode plots for ENET and SVMLIN and only one tree out of the 500 used in the RF model. Note: the features names are not readable due to the large number of variables in the model — here we focus on coefficient distribution).

BTR models can be illustrated as balances, where species abundance is symbolized by the cumulative weight (**Figure 1**). The concept of balances is not new in ecology and was first proposed to address the compositionality problem of microbiome data. A balance-based representation of the microbiome data can solve part of these issues and reveal biological patterns that were previously undiscovered ^27^. Very recently, other authors have applied the balance representation to the predictive context ^28^. Here, we propose more general models that encompass such balances (*i.e.* Ter models applied to log-transformed data - named TerLog; see supplementary material; **Figure S7**). Learning linear models on log-transformed counts correspond to identifying balances of multiplicative relationships. Which relationship best characterize microbial ecosystems remains an open question.

We devised a dedicated algorithm called *predomics* to learn BTR models from metagenomics data. Based on a genetic algorithm it supports learning high-quality models (see online methods). From a ML perspective, learning BTR models corresponds to minimizing the sum of a cost function (e.g. residual sum of squares (RSS)) and a L1 norm regularization for the sparsity, under a constraint on the unary value of the linear model that predicts classes.

### BTR models are accurate and improve with taxonomic specificity

Abundance can be quantified at different taxonomic levels. We generated BTR models on six different public metagenomic datasets (**Table S1**) and nine derived types of variables, (taxonomic levels, marker genes and pathway table, a fused taxonomic dataset, *i.e.* a total of 54 datasets, see online methods). We compared them with the SOTA algorithms: SVM, random forest (RF) and ENET. First, we tested models with different numbers of features (i.e. model-size) and noticed an effect on accuracy. Importantly, the testing performance was relatively different form training for the SOTA, indicating important overfitting. On the contrary, BTR models while being accurate, displayed comparable performance on both training and testing sets. The simplicity and sparsity of the BTR models diminishes overfitting (**Figure S1**).

We applied model-size penalization on the empirical accuracy to select the best model. All algorithms were evaluated by measuring test accuracy in a cross-validation setting and compared between them using paired tests (see online methods). BTR models performed at least as well as SOTA in 39/54 (72%). They outperformed SOTA in 1t/54 (30%) and were outperformed in 15/54 (28%) (**Figure 2; Figure S2A-C)**. RF displayed good results but at the expense of lower interpretability (hundreds of variables used in 500 trees; **Figure 2; Figure 3F**).

**Figure 2:**
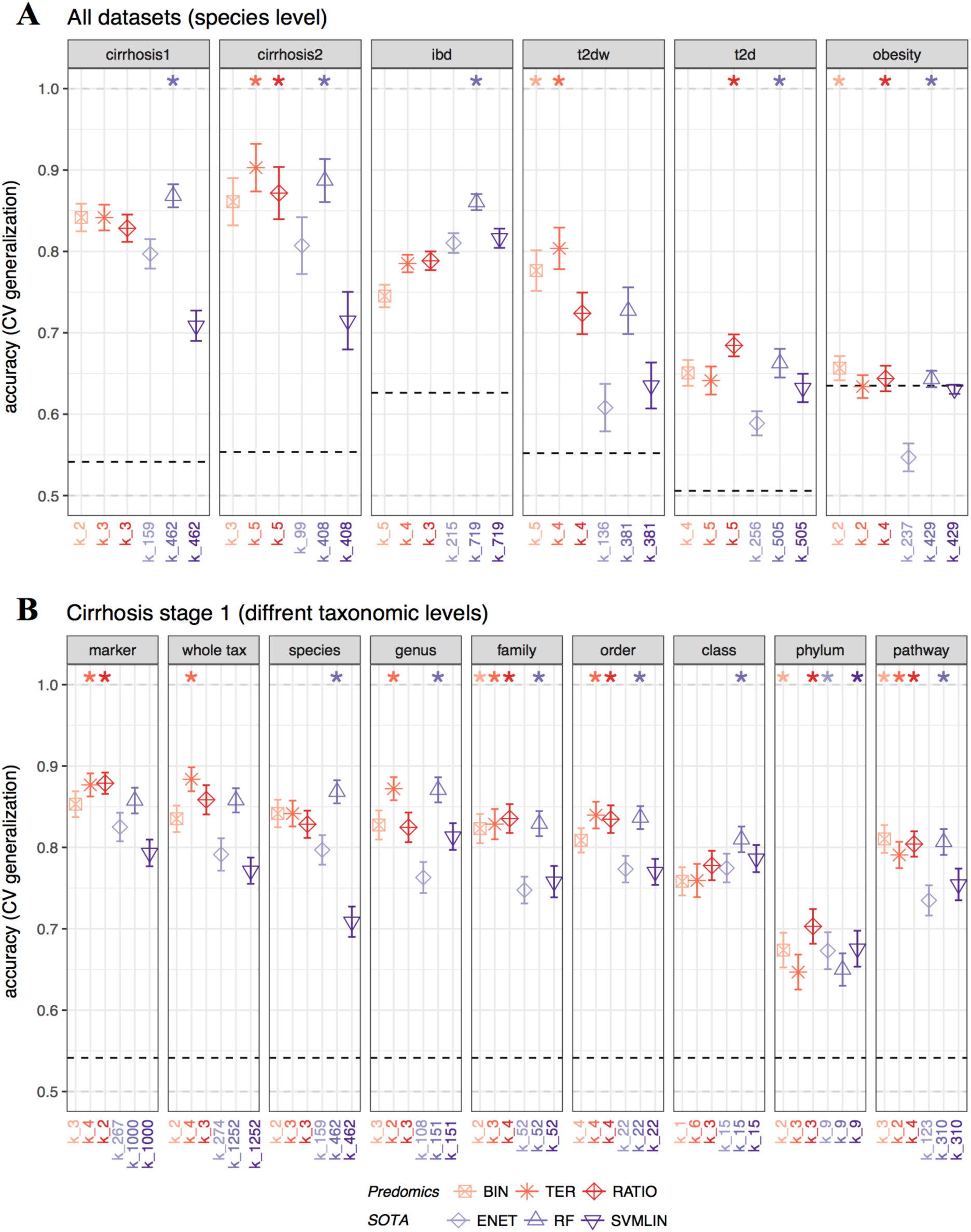
BTR models vs. SOTA performance across different disease and taxonomic levels. **A:** Accuracy measured in the test datasets at the species level across six different datasets. The stars on top indicate whether the corresponding BTR or SOTA algorithms are significantly better than others (*i.e.* without stars). **B**: Accuracy measured in the test datasets in different taxonomic levels of gut microbiome quantification at different phylogenetic levels (*marker gene, whole taxonomy, species, genus, family, order, class and phylum*). Dashed bars indicate the majority class and k_* indicates the model-size. A 10-times 10-fold validation test values are summarized as mean +/-standard errors.

Moreover, we tested the generalization of Bin, Ter, Ratio and TerLog models in an independent dataset. Learned in Cirrhosis stage-1, they were tested on Cirrhosis stage-2 dataset. Results illustrated in **Figure S4** indicate very good external validation with an average training accuracy=0.89 (sd=0.02) and testing accuracy=0.85 (sd=0.04). Ter and Ratio models generalized better compared to Bin and TerLog.

Results on different taxonomic levels (Cirrhosis stage-1) displayed higher performance at the gene marker, species and genus level, and decreased with higher taxonomic levels. Moreover, when applied to a multi-taxonomic level dataset (*strain* to *phylum* as generated by ^29^ with different specificity levels mixed together; *i.e. whole tax*), models displayed surprisingly good performance (**Figure 2B**). Indeed, in this space, models can be powerful as they can summarize more complex rules such as: “if abundance of *all Firmicutes minus all the Clostridiales order greater than threshold, then disease”*.

In addition to the abundance datasets described above and based on the zero-inflated nature of microbiome data, we trained and tested similar models on simple *presence* binary data derived from the previous 54 abundance datasets. Overall results are relatively similar indicating that the detection of species alone can be powerful in the prediction task (see supplementary material; **Figure S2D-F; Figure S3**). Noteworthy, when applied to presence data, BTR models indicate relationships between sub-ecosystem complexity or richness. These can be useful to detect switch-like mechanisms in the microbiome.

### BTR models are more interpretable than state of the art

A barcode graphical representation illustrates the simplicity of BTR models. In **Figure 3A-C** *left*, the models are represented by red and blue lines, corresponding respectively to positive and negative coefficients. Their length is proportional to the coefficient. The same representation is used to visualise the normalized coefficients of ENET and SVMLIN models, which include 159 and 4t2 variables respectively (**Figure 3D-E**). The RF model is more difficult to represent graphically and only one of the 500 decision-trees used in the model is illustrated (**Figure 3F**).

For each variable selected by BTR models, we assessed their importance in prediction. We implemented a variant of the well-known mean decrease accuracy (MDA) (**Figure 3A-C** *middle*; see online methods). Moreover, variable importance may differ from one model type to another. For instance, *Veillonella unclassified* is the most important for *Bin* and *Ter* but not for *Ratio*, which favours *Streptococcus anginosus*. Such importance score allows prioritizing further exploration of the features in the context of the predicted phenomenon.

*Predomics* can discover a family of BTR models with equivalent predictive power in a given model-size range (*i.e.* FBM for family of best models; **Figure S5**; see online methods and supplementary material). The selected FBM is analysed to identify the common features that are found in the models. For instance, in the cirrhosis stage-1 (species) dataset, the 2t8 models in the FBM with model-size<6 only rely on t7 features (i.e. 1t% of the whole dataset). This feature core-set, along with model copresence information allow us to infer a *model co-presence network*. It indicates the topology of information redundancy and complementarity in the predictive task (**Figure 4A**). In the left part of the network, there is a cluster of species that are more abundant/prevalent in the patient group and on the right, a cluster of species that are more abundant/prevalent in the healthy controls (**Figure 4B**). These species display negative intra-connectivity (inside modules) and positive inter-connectivity (among modules), indicating respectively exclusion and inclusion in the models. An emerging property of this network is the clustering of phylogenetically close species as depicted by node colours (blue tones for Firmicutes species enriched in patients and green tones for Proteobacteria and Actinobacteria enriched in controls), indicating information redundancy in prediction.

**Figure 4:**
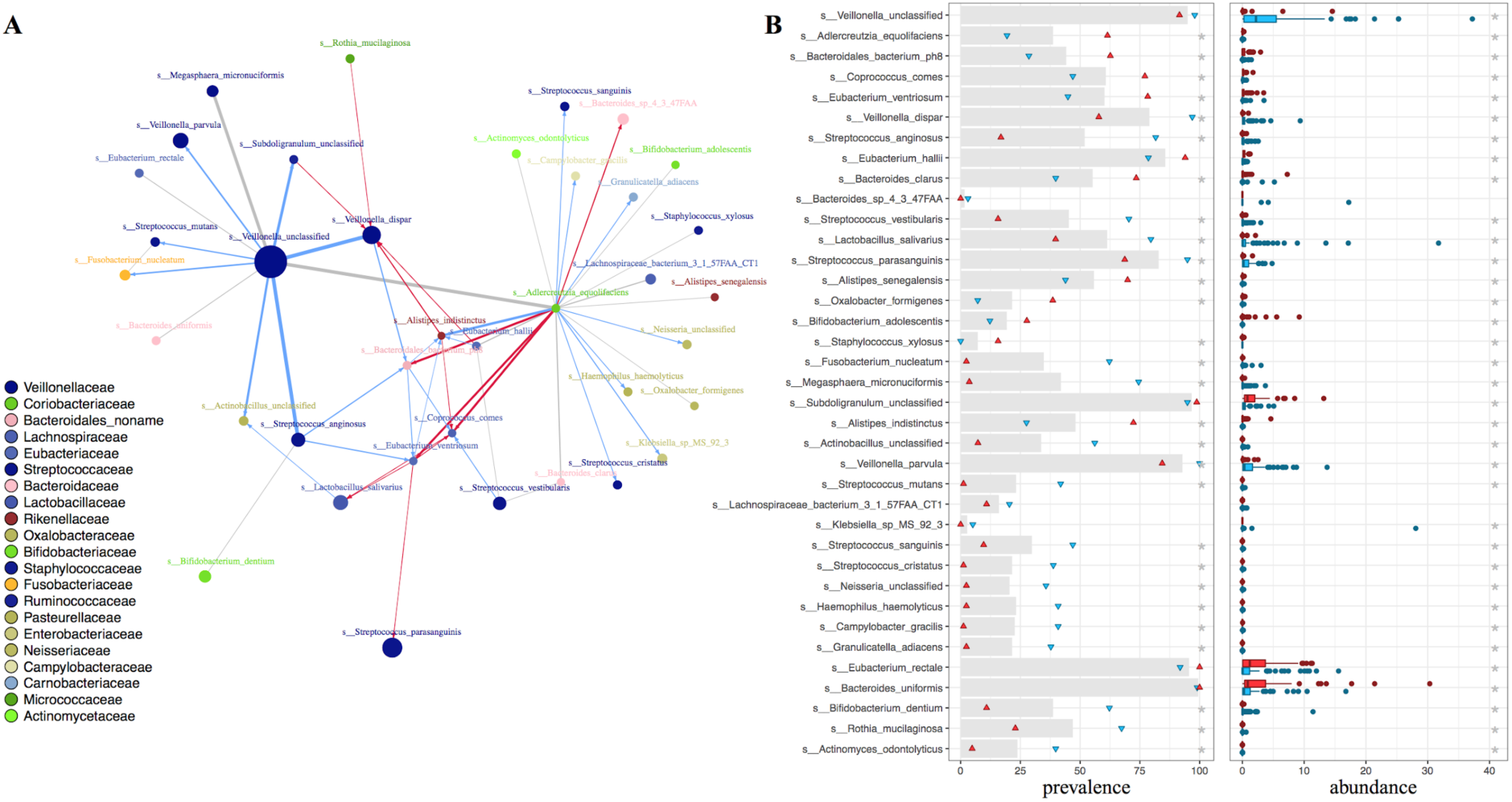
Information network of the most predictive FBM features. **A**: The network displays the top 5% strongest edges inferred using the ScaleNet network reconstruction approach (parameterized with *bayes_hc* and *aracne* methods, see online methods) in the FBM featurepresence table. The size of the nodes is proportional to the average importance (MDA) in the BIN, TER and RATIO experiments. The colours of the nodes indicate the taxonomic family assignation as indicated in the legend. The red and blue edges indicate co-presence and co-absence in the models respectively. **B**: For each feature present in the network we show in the *left*: the prevalence of the features in the whole dataset (grey bar) and in the prediction classes (disease, healthy) depicted as blue and red dots respectively and in the *right*: the feature abundance distribution in the prediction classes (disease, healthy) depicted as blue and red box plots respectively. Grey stars indicate significant differences.

Overall, BTR models, even those as simple as composed of five features or less, are surprisingly accurate and select important variables. Along with the corresponding visualisation interfaces and different statistics on feature importance (MDA; see supplementary material), these models are much more interpretable compared to state-of-the-art ones. Finally, by taking advantage of the FBM we provide useful insight in the predictive mechanisms of the variables and allow establishing confidence on the predictive core features as well as subsequent models (**Figure S5-S7**).

### BTR models provide biological insights

We evaluated BTR modelsp’ ability to provide biological insight in different medical conditions. We focused on the liver cirrhosis dataset ^7^, where major patient dysbiosis was observed with decreased microbial richness, depletion of gut commensals, and an invasion of oral bacteria. Several markers at taxonomic and functional levels were associated with the disease. *Predomics* BTR models replicated original findings (see supplementary material) and identified novel bacterial features associated with liver cirrhosis.

A more in-depth exploration of the species model co-occurrence network (**Figure 4**) emphasize the selection of strongly associated species as well as other redundant/complementary ones. As these species are phylogenetically close, they may offer similar functional services. Such is the case of an unknown species of *Veillonella* and *Veillonella dispar*. Notably, the model co-presence network, resembles the one described in the original study. The difference being that the original network was constructed using abundance of metagenomic species (i.e. MGS) for each metagenome. Thus, BTR models have the ability to distil and capture the biological information embedded in the data related to the prediction task.

Other authors have modelled liver cirrhosis associated microbiome using curated information from the literature, such as the ratio of autochthonous (butyrate-producer bacteria) to non-autochthonous (oral bacteria, opportunistic pathogens). The authors used these taxa to build a cirrhosis dysbiosis ratio (CDR) score ^30^. Based on their description we built three redundant models using family taxonomic features and tested them in the liver cirrhosis stage 1 (family) dataset ^7^. The *predomics* Ratio model provided far superior performance (accuracy=0.86) compared with CDR-based models (accuracy=0.56 in average) (**Figure 5**). The reason for CDR lower performance can be explained by the inclusion of the *Bacteroidaceae* family in the liver cirrhosis group. However, we observe the opposite association in the current dataset, where *Bacteroidetes*-related features are enriched in the control group. This is consistent for different taxonomic levels (**Figure 5**, see supplementary material).

**Figure 5:**
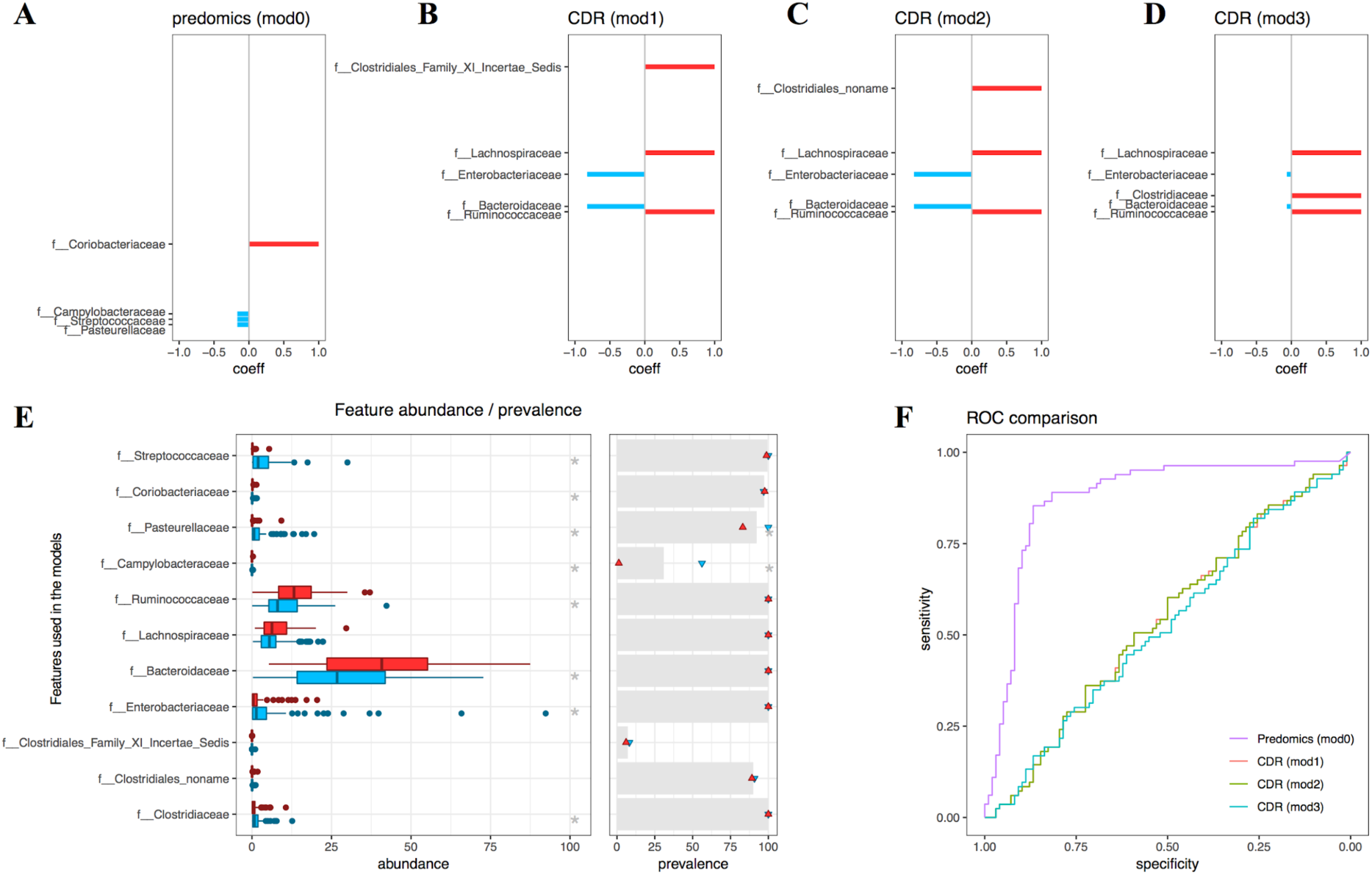
Cirrhosis Dysbiosis Ratio (CDR) index compared to *predomics* ratio model. **A-D**: Barcode plots indicating the coefficients of the Ratio models *(S13-S15)* build with features from the CDR index and *predomics* discovered model *(S16)*. Red and blue colours indicate respectively the numerator and denominator of the ratio model and are respectively enriched in the controls and LC patients. The length of the lines is proportional to the ratio factor optimized in the model. **E** *left*: Boxplots indicating the abundance distribution by class for all features used in these models (red is enriched in controls and blue in the liver cirrhosis group). *right*: for the same features the prevalence of non-zero values is depicted in grey for the whole cohort and red and blue dots respectively in the control and patient groups. Grey stars indicate significant difference. **F**: Receiver operating characteristic (ROC) curves for the four models *(S13-S1t)*.

**Figure 6:**
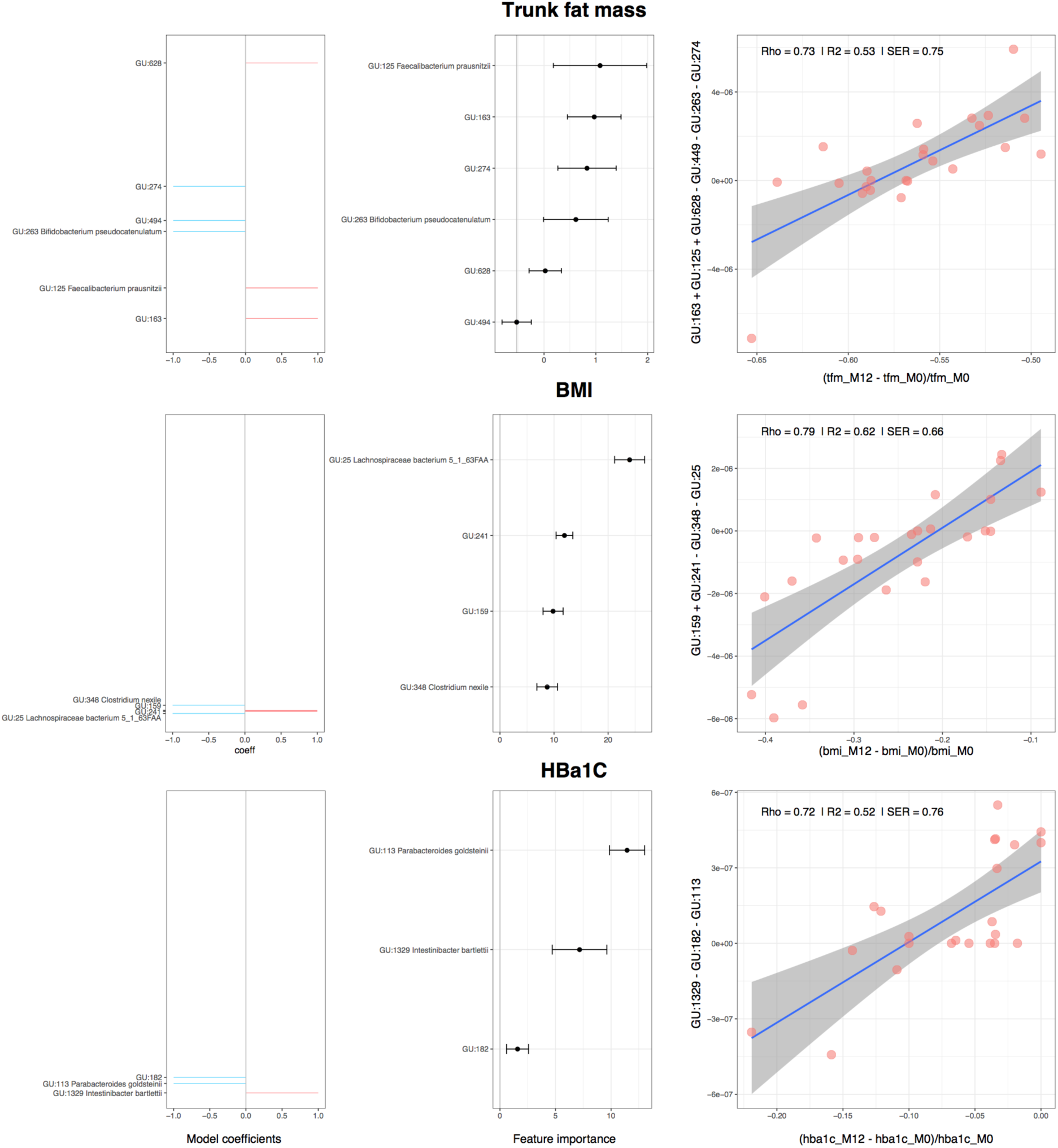
Quantitative prediction of phenotypic outcome after bypass surgery. *Left*: Barcode plots indicating the coefficients of the ternary models. *Middle*: Percentage of mean decrease R^2^, measuring the importance of features in the fitting objective during the cross-validation process. *Right*: Scatter plots indicating the fitting of the model score (y-axis) as measured with baseline microbial profiles against the relative change of each of the three phenotypes (trunk fat mass, BMI and HbA1C) in the x-axis.

### Quantitative prediction using BTR regression models

In addition to classification, *predomics* can perform regression tasks by searching models that correlate with the quantitative variable to predict (see online methods). We used data from a recently published study where obese patients underwent Roux-en-Y gastric bypass (RYGB; n=14) and adjustable gastric band (AGB; n=10) surgery ^31^. Patientsp metagenomes were measured pre-surgery and twelve months post-surgery (among others). Most patients who underwent the surgery improved their body weight, body composition and glucose homeostasis (glycemia, insulinemia and glycated haemoglobin (*i.e.* HbA1C)) with significant variation between individuals. Metabolic improvement was measured as the relative change at 12 months compared to baseline.

We searched pre-surgery metagenomic data for bacteria that could predict the improvement of BMI, trunk fat distribution, and HbA1C and discovered models composed of t, 4, and 3 species reaching R^2^ values of 0.53, 0.t2, 0.52 respectively (**Figure t**). The algorithm generalizes well when tested in cross validation (20-times 5-fold CV), although we observe decreasing performance in testing sets likely due to the small sample size. Interestingly, the models highlight bacterial species such as *Faecalibacterium prausnitzi, B. pseudocatenulatum and P. goldsteni*, which were previously shown to be associated with metabolic health and low-grade inflammation (see supplementary material). While this is a proof of concept, these results illustrate the power of the microbiome to predict change in body composition and glucose homeostasis.

## DISCUSSION

In principle, BTR models could be applied to any type of data. However, they are best suited to commensurable measurements (*i.e.* variables measurable by the same standard or measure). In the growing field of metagenomics, issues related to compositionality and data processing still remain to be solved. Recent work has shown the importance of data acquisition in subsequent analytical inferences. In particular, microbial loads differ significantly between individuals and are associated with specific types of microbial ecosystems ^32^. An advantage of *Ratio* models is that they are scale-invariant given they do not depend on absolute measurements, thus avoiding compositionality issues.

Algorithmically, the complexity of finding optimal BTR models grows exponentially with the number of variables. However, it has been proven that near-optimal binary-weighted models can be identified in polynomial time ^33^. In *predomics*, we implemented several heuristics that support finding near-optimal models while remaining scalable (see online methods).

The simplicity of a BTR model may come with the risk of over-interpretation. The existence of *k* species in a model, may correspond to different explanations ranging from simple correlation to causal relation. They may or may not interact together, as in the case of a niche differentiation ^28^. For instance, the buccal-originated species found in the gut of liver cirrhosis patients ^7^ along with the absence of commensals may reflect a global difference in the environment where they live rather than direct interaction ^7^. Even if BTR models represent real interactions between species, it is not recommended to give a causal interpretation without experimental or literature validation. Nevertheless, identifying such species provides important knowledge towards understanding potential mechanisms between species or between species and the host.

The quality of reference datasets used in building predictive models is vital for model interpretability. The propagation of errors and inaccuracies in genomic datasets is a well-known issue, and affects automated methods for functional annotation ^34^. Moreover, due to the lack of biochemical characterization of orphan enzymatic activities, the number of sequences with unknown functions is extremely large, making their propagation common ^35^ (see supplementary material).

Another caveat with microbiome studies resides in the potential confounders modulating microbial ecosystems. For instance, metformin can alter the bacterial ecosystem such that some bacterial species (*e.g. E. coli*) are increased in abundance and others are depleted ^36^. It is important to filter out confounder-related species from the data or to filter out models that are sensitive to confounders.

Finally, after filtering for confounders, and manual curation, BTR models can be used to develop specific acquisition technologies such as microarray DNA chips, built with primers that are specific to the species found in the models ^37^. From a clinical perspective, identifying a small subset of variables (genes, species, pathways, etc) can be used to simultaneously predict multiple tasks. Such applications, after being properly validated, will be important to the medical community in their translational quest to improve patient care. Our *predomics* approach brings us a step closer towards this goal.

## METHODS

Methods and any associated references are available in the online version of the paper. Supplementary information and source data files are available in the online version of the paper. The *predomics* package is available in *https://git.integromics.fr/published/predomics*.

## ACKNOWLEDGEMENTS

This work was made possible by the Funding Support of European Unionps’ Seventh Framework Program under grant agreement HEALTH-F4-2012-305312, by the Assistance Publique-Hôpitaux de Paris promoter of the clinic program. This work was also supported by the French National Agency through the national program Investissements d’ Avenir (reference no. ANR-10-IAHU-05) IHU ICAN. By Assistance Publique-Hôpitaux de Paris Contrat dpinterface chercheurs 2015-2018. We wish to thank S.D. Ehrlich as well E. Le Chatelier for mindful discussions on the early stages of this work and T. Swartz for help in language proofreading.

## AUTHOR CONTRIBUTIONS

**Prifti.E**: overall conception, design and interpretation; designing and coding the software; conducting all experiments; writing the manuscript; final approval of the manuscript.

**Chevaleyre.Y** conception and interpretation of the approach; coding; drafting, approving the manuscript.

**Hanczar.B**: conception, design and interpretation of the approach; coding; drafting, approving the manuscript.

**Belda.E**: biological interpretation of the results; drafting, approving the manuscript.

**Danchin.A**: biological interpretation of the results; drafting, approving the manuscript.

**Clément.C**: data production (bariatric model); biological interpretation of results; approving the manuscript.

**Zucker.J-D**: conception, early prototyping, design and interpretation of results; drafting and final approval of the manuscript.

## COMPTETING FINANCIAL INTERESTS

The authors declare no competing financial interests.

## ONLINE METHODS

### Public datasets used in this study

To test *predomics* and compare it with state-of-the-art methods, we used several public datasets. For the classification tasks we downloaded five curated metagenomic datasets from the ExperimentHub ^29^. They were generated in several independent studies using shotgun metagenomics (**Table S1**) and were processed bioinformatically and curated independently by Pasolli et al ^29^. In short, metagenomes were sequenced on the Illumina platform at an average depth of 45 Million reads. MetaPhlAn (v2.0) was used on the pre-processed reads with default settings to generate microbial community profiles (from kingdom to species taxonomic levels). To obtain functional profiles, HUMAnN23 (v0.7.1) was used on the pre-processed reads with default parameters. Three main outputs: gene family abundance, pathway abundance, and pathway coverage were generated. The R code used to download and format the data used in our experiments is provided in the supplementary material package. We derived 54 datasets out of the original data (i.e. six different cohorts and for each six taxonomic levels, a marker gene and a pathway table along with a fused taxonomic dataset). These same 54 datasets were also transformed as presence/absence derivatives used for additional experiments

For the regression experiments, we used shotgun metagenomics data from a recently published study, where morbidly obese patients’ underwent bariatric surgery ^31^. Patientsp microbial DNA was sequenced using SOLiD before surgery and one, three, and twelve months after surgery. Reads were cleaned and contaminants were removed before mapping them against the 3.9M gene catalogue ^19^. Counts were rarefied at 11 million reads and normalized. Metagenomic species (MGS) abundance was computed as the average of the 50 most connected genes of each MGS after 20% presence filtering ^19^. See original study for more information ^31^.

### Novel ecologically inspired models

The new family of BTR models (for Binary/Ternary/Ratio) is inspired by possible relationships between species within an ecosystem ^13^ with *X*_1_… *X*_*p*_ the predictor variables of a metagenomic sample. For simplicity, *X*_*j*_ represents the abundance of the *j*^*th*^ bacterial species, and each patient can be classified into two conditions: healthy or diseased. Until now, the algorithms yielding the most interpretable models are based on sparse logistic regression such as the Lasso algorithm ^23^ or its improvement the Elastic Net algorithm. Here, we argue that it is possible to consider even simpler models that improve interpretability without worsening the accuracy when compared to state-of-the-art algorithms.

The implicit biological assumptions underlying the explicability of such linear models are: 1) the contribution of each bacterial species to the prediction is proportional to its abundance, and 2) that only a limited number of species are sufficient to support the prediction. Our BTR models are inspired by three hypotheses emphasizing relationships between species and associated ecosystemic outcomes. They are particular types of linear models, which are generally a sequence of real coefficients *β*_0_ *β*_*p*_. A patient is predicted as ill with a probability *p* > 1/2 if and only if 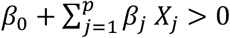

The *binary models* (*i.e.* Bin) are defined based on the first hypothesis that the unweighted cumulative abundance of a limited group of species may be sufficient to support the prediction. This translates in a linear model with the additional constraint that each coefficient *β*_1_… *β*_*p*_ (omitting the intercept *β*_0._) is limited to the value 0 or 1. An example of a binary model is in *(1)*, **Figure 3A**, which may be interpreted as “if the cumulated abundance of *s Veillonella_unclassified* and *s Clostridium_perfringens* is smaller than 0.18 (i.e. 18% of the total microbial abundance), then the individual is classified as healthy”. Such model can correspond to the end result of different types of relations: either no direct interaction between these species (e.g. use non-overlapping resources of the corresponding environment or not colocated), or a real interaction (be it cooperation or competition as both are possible) ^25,26^.

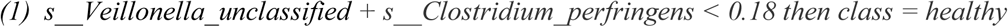

The *ternary models* (*i.e.* Ter) are defined based on the second hypothesis that both cumulative and difference of abundance of a limited group of species may be enough to support the prediction. This translates in a linear model with the additional constraint that coefficients *β*_1_… *β*_*p*_ (omitting the intercept *β*_0._) are limited to the value −1, 0 or 1. An example of a ternary model is in *(2)*, **Figure 3B**. It may be interpreted as follows: “if the abundance of *s Alistipes_indistinctus* minus the cumulative abundance of *s Streptococcus_anginosus* and *s Veillonella_unclassified* is greater than −0.083, then the patient is classified as being healthy”. Such model can correspond to the end result of different types of interactions including cooperation between *s Streptococcus_anginosus* and *s Veillonella_unclassified* and also competition between both species and *s Alistipes_indistinctus*. For Bin and Ter models we can optionally constrain the intercept to be equal to zero.

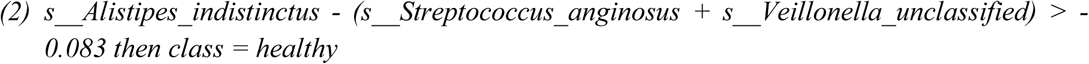

Finally, the *ratio models* are defined based on the third assumption that the disease state of the patient may be determined by the ratio of the cumulative abundance of two groups of species rather than their difference. These are also linear models with an additional constraint: each coefficient *β*_1_… *β*_*p*_ is constrained to have either the value *-θ*, 0 or 1, where *θ*a positive real number, and the intercept, *β*_0_, is set to zero. An example of a ratio model is in *(3)*, **Figure 3C**. It may be interpreted as follows: “if the abundance of *s Subdoligranulum_unclassified* is *θ* = 81 times greater than the total abundance of the group of species *s Megasphaera_micronuciformis + s Streptococcus_anginosus* then the individual is classified as healthy”.

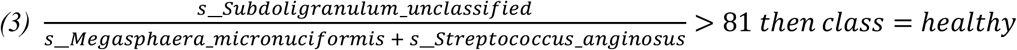

BTR models can be used for classification regression tasks. In this case once the modelps score is computed, two additional parameters are learned *α* and *β*_.0_ using a linear regression to estimate a transfer function from the microbiome relative abundance to that in which the predicted variable is expressed.

### The predomics optimization algorithm

Learning optimal BTR models is a very hard computational task. Because weights are discrete, usual techniques coming from convex optimization do not apply here. A naïve way to generate optimal BTR models would be to perform an exhaustive search though the space of all models. Unfortunately, this is not feasible in practice because the computation time would be exponential with the number of features. The BTR learning problem is known as NP-Hard (a notion from computation complexity theory), which means that *no* algorithm can solve this problem exactly in polynomial time, unless a widely believed conjecture turns out to be false ^33^.

Because tractable optimal algorithms are out of reach at the moment, we can only apply approximate heuristic methods, without any guarantee on the optimality of the outcome. One such popular family of heuristics is the genetic algorithm, which is a stochastic optimization technique. It adopts concepts from evolutionary biology — populations, reproduction, mutation and generations. Although the general principle behind all genetic algorithms is the same, the strategies used can be tailored and so did we for the problem of BTR model construction. The outline of the algorithm is described below, and the full implementation is provided along with the supplementary material package.

1. The first step of any genetic algorithm is the generation of an initial set of candidate models, which is usually called the “initial population” *p*_*t*_, typically composed of 100 random models. This step is crucial, because it sets the initial exploration space. Most existing genetic algorithm build this set by simply drawing random candidate models. In our case, we observed that combining various algorithms to generate this initial population boosts the final accuracy. More precisely, we combine models generated by a beam-search algorithm, models obtained by a logistic regression followed by a weight discretization phase (^33^) and purely random models. These models are chosen of different sizes (i.e. parsimony), typically ∈{1: 30}.

2. Then, the algorithm performs 100 iterations (if no convergence criterion is set). At each iteration *t*, the algorithm generates a new set of improved models *p*_*t*+1_(a new population) based on the previous population *p*_*t*_. More precisely, to build *p*_t+1_ the algorithm performs four consecutive stages, which are the *evaluation, selection, crossover* and *mutation*.

a. During the evaluation stage, all models in *p*_*t*_ are evaluated according to their predictive accuracy. Each of the three remaining stages outputs a modified population based on the population of the previous stage.
b. Typically, 50% of the models are selected half randomly and half based on the best accuracies. This selection will be at the origin of the new generation of models *p*_*t*+1_.
c. During the cross-over stage, pairs of models are randomly drawn among those who survived the selection stage, and their features are combined randomly to generate new models, which are added to the population.
d. In the mutation stage, a fraction of the models, selected randomly, are mutated. The mutation of a candidate model is the process of either removing, or adding a random feature, or even altering the weights of one or more of its features. At this stage the *p*_t+1_is created and will serve as initial population of *p* _t+2_ and so on.

3. At the end of the evolution process, a population of models *p* _*final*_ is provided on which the family of best models (FBM) can be selected. The best model is obtained by applying a model-size penalization. The penalized accuracy *accuracy*_*penalized*_ = *accuracy* - *ƛ k* is computed, where *k* is the number of features in the model (i.e. parsimony) and *ƛ* is an hyperparameter controlling the penalization of the accuracy. The number of selected features in the BTR models depends on this hyper-parameter and will increase when *ƛ* decreases. We believe that applying the sparsity bet criterion to our models will improve their overall generalization.

For classification tasks, we can optimize different parameters such as accuracy (default), AUC, F1, precision or recall. For regression tasks we can optimize the rho, R^2^ (default) or standard error of the regression. The interested reader may look at our code for more information, available at the projectps repository https://git.integromics.fr/published/predomics.

### Experimental design

The BTR models are tested on the 109 different datasets (see above) and compared with the methods from the state-of-the-art machine learning algorithms: support vector machine (SVM) with linear and Gaussian kernel (data not shown), Random Forest and Elastic Net (an improvement of Lasso, alpha=0.5). A more specific comparison between TerLog models and the geometric mean balance algorithm is provided in supplementary material. Our experimental pipeline proceeds as follows:

1 Feature normalization: frequency tables are used as already processed by Pasolli et al ^29^.

2.Features with low standard deviation are filtered out. The threshold corresponds to the maximum second derivative of the distribution of the featureps standard deviation.

3 The accuracy in generalisation of each method is estimated by 10-times 10-fold cross validation for the classification tasks and a 20-times 5-fold cross validation for the regression tasks. Each model is fitted on the training data of each cross-validation fold and tested on the corresponding test data.

4.The feature selection is embedded for the BTR models and Elastic Net. For SVM and random forest we do not apply a feature selection, besides the model size experiments (**Figure S1**) where feature selection is based on the Mann-Whitney score test score as introduced in ^38^. The feature selection step is included in the cross-validation loop in order to avoid the selection bias^39^.

5.Once the best learner is identified, it is compared with a paired t-test with all others over the 100 CV points. Those that are not significantly different are considered equivalent (pval<0.05).

### Family of best models

A family of best models (i.e. FBM) is defined as the set of models returned by the *predomics* algorithm, whose accuracy is within a given window of the best model accuracy. This window is defined by computing a significance threshold assuming that the model accuracy follows a binomial distribution (p<0.05). An FBM can be analysed in detail to distil biological information in the predictive context (see supplementary material).

### Feature importance

Similarly, to what is proposed in the random forest algorithm, feature importance is defined as the usefulness of features to predict, given all other features and best models of the FBM. The first step is to perform a k-fold cross-validation of the learning algorithm. During each fold, the out-of-bag error on each model composing the FBM is computed. To measure the importance of the *jth* feature, its values are permuted within the out-of-bag data and the out-of-bag error is again computed on this perturbed data set for each FBM model. The *importance score* for the *jth* feature is computed by averaging over all FBM models the difference in out-of-bag error before and after the permutation. The mean decrease accuracy (MDA) is finally computed as the average of these values over all the folds and is displayed along with the standard error of the mean.

Moreover, we propose a second concept of importance (named PDA), based on the mean prevalence in FBM models (i.e. percentage of times a given feature is selected in a model composing the FBM). Next, we compute the average model prevalence for each fold during the cross-validation process and finally, propose the CV-averaged PDA along with the standard error of the mean as measure of feature importance. This is illustrated in **Figure S8B**.

### Regression models

*Predomics* can learn also regression models. These models are evaluated by either maximizing, Spearman rho or Spearman R^2^ or minimizing the scaled standard error of regression (SER). The model score at this stage reflects the cumulative/difference/ratio of relative abundance of the species and needs to be scaled in the ranges of the variable to predict. For this reason, in this regression setting, we estimate two additional parameters alpha (i.e. multiplication factor) and beta (i.e. intercept).

### Network reconstruction

Different methods can be used to reconstruct the feature co-presence network in model selection data. Here we used the Scalenet methodo ^40^. To accurately reconstruct such network ScaleNet first reduces the reconstruction problem into a large number of simpler reconstruction problems, then employs state-of-the-art reconstruction methods to solve them. Finally, a consensual voting strategy between the methods is adopted to identify the most accurate sub-graphs. The different sub-graphs are then connected like pieces of a larger puzzle. The main originality of the method lies in its powerful problem reduction based on spectral decomposition. Learning small problems instead of a large one with few observations as is the cas in omics data is showed to lower the overfitting effect ^40^. Here we used the top 5% strongest edges inferred using the ScaleNet network reconstruction approach (parameterized with *bayes_hc* and *aracne* methods) in the FBM-presence table.

